# Inorganic sulfate transport by the *Mycobacterium tuberculosis* PE22/PPE36 complex

**DOI:** 10.64898/2026.06.04.730267

**Authors:** Ashutosh Tripathi, Vishant Boradia, Amy Wu, Marguerite Dawkins, Anas Saleh, Kyu Y. Rhee, Christoph Grundner

## Abstract

*Mycobacterium tuberculosis* (*Mtb*) encodes two multigene families with 169 members that are exclusive to mycobacteria, the *pe* and *ppe* genes. These genes have unusual sequences including low-complexity repeat regions, but their functions—and whether they share a common function—have long been unclear. Recently, several members of the *pe/ppe* family were shown to transport nutrients across the outer *Mtb* membrane, a role for which no other proteins have yet been identified. Whether nutrient transport is a family-wide function and the range of nutrients transported by the PE/PPEs remains unclear. Sulfur is an essential nutrient for *Mtb* physiology and pathogenesis. To test whether PE/PPE transporters contribute to sulfur acquisition, we analyzed the transcriptional response of *Mtb* to sulfate by RNA sequencing. The *pe22/ppe36* genes were induced in sulfate-limiting conditions. Deletion of *pe22/ppe36* impaired growth in low-sulfate media and reduced intracellular sulfate levels, effects that were reversed by heterologous expression of the *Mycobacterium smegmatis* porin MspA. The response of sulfur-responsive genes to sulfur was muted in the *pe22/ppe36* deletion strain, and mass spectrometry showed lower sulfolipid and sulfur metabolite levels in the deletion strain. These findings identify PE22/PPE36 as a specific sulfate uptake system and supports the emerging idea of PE/PPE proteins as nutrient uptake systems across the *Mtb* outer membrane.

**Significance statement:** The mechanism of nutrient transport across the porin-less mycobacterial outer membrane and the function of the large *pe/ppe* gene family have been longstanding questions in mycobacterial biology. This study shows that the *Mycobacterium tuberculosis* PE22/PPE36 complex serves as a selective conduit for inorganic sulfate. Deletion of this complex disrupts intracellular sulfur homeostasis and triggers a metabolic seesaw that depletes cell-surface sulfolipids. Our findings expand the PE/PPE transport paradigm and suggest a direct link between nutrient acquisition and TB transmission.

## Introduction

*Mycobacterium tuberculosis* (*Mtb*) is surrounded by two distinct membranes: the plasma membrane and the outer, or mycomembrane (1). The outer membrane is covalently linked to an arabinogalactan–peptidoglycan complex and contains diverse noncovalently associated lipids such as phthiocerol dimycocerosate (PDIM) and sulfated trehalose glycolipids (SGLs) (1). Unlike Gram-negative bacteria, however, *Mtb* does not express recognizable porins to allow nutrients across the outer membrane. As a result, *Mtb* is approximately 100- to 1000-fold less permeable to small hydrophilic solutes than Gram-negative bacteria such as *Escherichia coli* and *Pseudomonas aeruginosa* (2). How nutrients cross the mycomembrane despite its extreme impermeability and lack of porins has been a longstanding question.

While *Mtb* and other slow-growing or pathogenic mycobacterial species lack canonical porins, this lack of porins correlates with an expansion of the PE/PPE protein families, suggesting that PE/PPE proteins may serve as functional analogs of porins. Supporting this idea, several PE/PPE proteins localize to the outer membrane, are secreted via the ESX (type VII) secretion systems, and have indeed recently been implicated in the transport of solutes such as Ca²⁺ (3) glucose, and glycerol (4). Also, PPE36 has been shown to be involved in heme acquisition (5), and several others in the transport of drugs (6). These observations collectively point toward a broader role of these unusual proteins as outer membrane transporters in *Mtb* and suggest a large system of specific, idiosyncratic transporters that functionally replace porins.

Inorganic sulfate (SO_₄_²⁻) is an essential *Mtb* nutrient required for the synthesis of sulfur-containing amino acids, iron–sulfur clusters of enzymes, redox buffers, and sulfolipids—key components of the mycobacterial cell envelope and important virulence factors (7). *Mtb* can metabolize organic sulfur-containing molecules but also uses inorganic sulfate as a source of sulfur. A recent study showed that inorganic sulfate is in fact the primary sulfur source of *Mtb in vivo* (8): Intracellular *Mtb* accumulated nearly 100-fold more inorganic sulfate than its host macrophages, reflecting a strong sulfur demand during infection that is primarily satisfied by inorganic sulfate. Free sulfate in the cytoplasm is tightly regulated as excessive reducing equivalents are toxic (9) and sulfate depletion triggers compensatory recovery through reverse transsulfuration pathways (10). The requirement for calibrated sulfate levels implies the existence of a dedicated sulfate transport system in the mycobacterial envelope. While the inner membrane ATP-binding cassette (ABC) transporter complex SubI–CysTWA mediates sulfate translocation across the inner membrane (11, 12), the mechanism by which sulfate crosses the outer membrane of *Mtb* is unknown.

Here, we sought to identify how *Mtb* acquires sulfate across the *Mtb* outer membrane. Transcriptomic analysis of *Mtb* under sulfate-replete and -depleted conditions showed upregulation of the *pe22* and *ppe36* genes. Genetic disruption of the *pe22/ppe36* operon resulted in growth defects under sulfate-limiting conditions and a reduction in intracellular sulfate accumulation, which was restored by genetic complementation or expression of the heterologous porin MspA. The *pe22/ppe36* knockout also had reduced sulfolipid levels. These findings reveal a previously uncharacterized sulfate transport system in *Mtb* and support the emerging idea that PE/PPE proteins are outer membrane transporters.

## Results

### Sulfate induces expression of *pe/ppe* genes

Sulfate plays a central role in *Mycobacterium tuberculosis* (*Mtb*) physiology, and recent studies have shown that inorganic sulfate is the primary source of sulfur for intracellular *Mtb* (8). While sulfate crosses the inner membrane through the SubI–CysTWA ABC transporter, it is not known how sulfate crosses the highly impermeable outer membrane. Transporters often respond transcriptionally to their substrates, which has also held true for some outer membrane transporters in *Mtb* (3, 13, 14). To understand the transcriptional response of *Mtb* to sulfate and identify candidate outer membrane transporter genes, we carried out an RNA-seq experiment to compare *Mtb* grown without sulfate and with 100 µM Na₂SO₄ supplementation.

We observed significant downregulation of *cysT, cysW, cysA2,* and *cysA3*, genes associated with the SubI–CysTWA ABC transporter operon also known as sulfate activating operon (SAC operon) in the sulfate-repleted condition after 48 hours (Fig. 1 A and B); another example of transporters responding transcriptionally to their cargo. Genes involved in cysteine biosynthesis (*cysK1, cysK2*) (Fig. 1 B) and components of the antioxidant machinery (*ahpC, ahpD, trxC*, and *thiS*) were also downregulated in the presence of sulfate (Fig. 1 A). Conversely, their upregulation under sulfate-deplete conditions suggests a response to elevated reactive oxygen species (ROS) as biosynthesis of cysteine-mediated redox buffer molecules (mycothiol, ergothiol and gammaglutamylcysteine) and iron-sulfur clusters necessary for maintaining redox balance are impaired (8, 15–17). In addition to these genes previously implicated in sulfate transport or metabolism, we identified 17 *pe/ppe* family genes with significant differential expression (Fig. 1C and Table 1). Among these, the co-operonic heterodimeric partners (18) *pe22* and *ppe36* were the most strongly upregulated, with ∼3-fold increase in transcript in response to sulfate.

**Fig 1.**
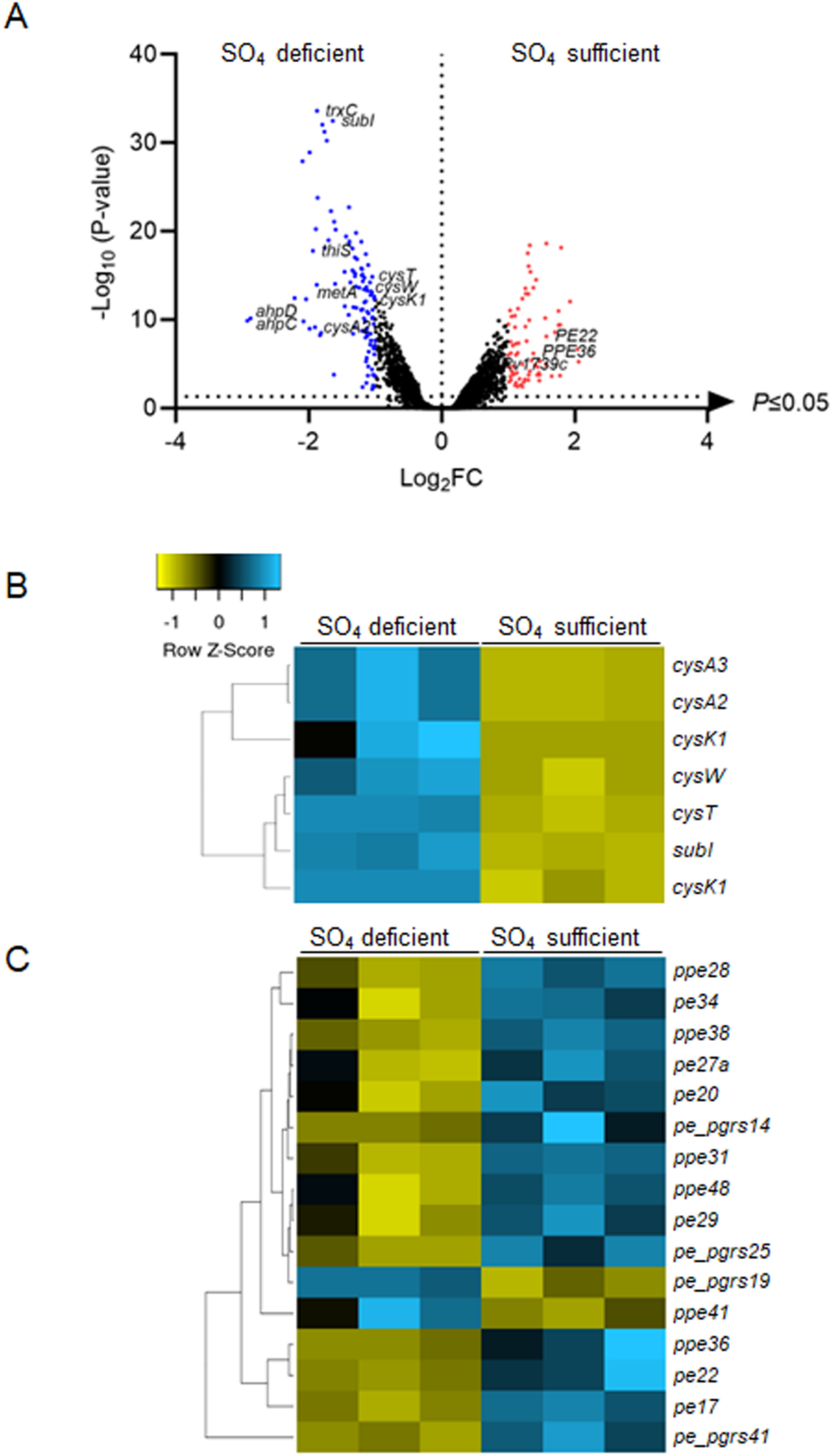
Sulfate downregulates inner membrane sulfur transporter genes and induces expression of *pe22/ppe36* genes. (A) Effect of sulfate on the WT *Mtb* transcriptome. Volcano plot shows log₂ fold-changes in gene expression under sulfate-replete and sulfate-deplete conditions (x-axis). The horizontal dotted line indicates the significance cutoff (P < 0.005). Significantly upregulated (≥1.5-fold) and downregulated (≤ 0.5-fold) genes are shown as red and blue dots, respectively, under sulfate-replete conditions. (B) Heat map showing sulfate transporter genes from the inner membrane sulfur ABC transporter and cysteine synthesis pathways. (C) Heat map showing significantly upregulated *pe/ppe* genes under sulfate-replete conditions. Significance was determined by one-way ANOVA.

**Table 1:** *Mycobacterium tuberculosis pe/ppe* genes upregulated after 48 h under sulfate-sufficient conditions. *: housekeeping control. ns: not significant.

|  | Gene name | Fold change | Avg *P*-value |
| --- | --- | --- | --- |
| 1 | *pe22* | 2.98 | 7.88E-09 |
| 2 | *ppe36* | 2.60 | 6.57E-07 |
| 3 | *ppe28* | 2.32 | 1.10E-05 |
| 4 | *pe29* | 1.95 | 1.41E-03 |
| 5 | *pe34* | 1.94 | 5.94E-03 |
| 6 | *ppe31* | 1.93 | 9.43E-06 |
| 7 | *ppe38* | 1.91 | 7.55E-08 |
| 8 | *pe27a* | 1.90 | 5.65E-03 |
| 9 | *sigA* * | 1.05 | ns |

### Depletion of *pe22/ppe36* results in reduced growth during sulfate limitation

Because *pe22* and *ppe36* are co-operonic and a known functional *pe/ppe* pair that forms a heterodimer, and because it showed the strongest response to sulfate, we chose *pe22/ppe36* for further study. Sulfur is essential for the growth of *Mtb* as demonstrated by the absence of growth for WT strains under sulfur-depleted conditions (Fig. 2A); however, sulfur-containing reducing equivalents such as cysteine, as well as the non-reducing derivative 3′-phosphoadenosine-5′-phosphosulfate (PAPS), are toxic to *Mtb* when present at high concentrations (7, 17). Together, these observations suggest that *Mtb* must maintain sulfur at a tightly regulated physiological concentration in the cytoplasm, making outer-membrane sulfur transporters a critical determinant of sulfur homeostasis.To test a potential role of *pe22/ppe36* in sulfate transport, we generated a *pe22/ppe36* knockout (KO) strain by recombineering (19) as well as a complemented strain (Comp) by introducing an episomal copy of the *pe22/ppe36* coding region under control of an inducible promoter into the *pe22/ppe36* KO strain. We tested the growth of the *pe22/ppe36* KO strain and WT in a range of inorganic sulfate concentrations (1mM–0.008mM). The sulfate concentration within the phagosome is not known. However, in contrast to the relatively abundant sulfate levels in human plasma (20), RNA-seq data of intracellular *Mtb* within macrophages (21), suggest that the pathogen experiences sulfate scarcity in the nutrient-deprived phagosomal environment of infected macrophages. Thus, we included sulfate concentrations well below the level in human plasma in our assay. *Mtb* strains were grown in sulfate-free medium for 48 hours, followed by sulfate repletion. After three weeks of sulfate repletion, none of the strains grew without sulfate, confirming that sulfate is essential for growth (Fig. 2A). When grown in low sulfate, the *pe22/ppe36* KO strain showed reduced growth in a spotting assay on 7H10 plates compared to the WT strain. Complementation of *pe22/ppe36* restored growth to WT levels, even at the lowest sulfate concentration tested (0.008 mM, which did not support growth of the WT). To quantify these growth differences, we plated WT and mutant strains grown in the same sulfate concentrations on solid agar for enumeration of colony forming units (CFU). Consistent with the spotting data, growth of the *pe22/ppe36* KO strain was severely impaired, with ∼3 log less CFU at a sulfate concentration at which WT grew normally, and no detectable CFU at lower concentrations. Complementation reverted these growth defects to WT. Interestingly, in the lowest sulfate concentration tested, WT and *pe22/ppe36* mutant were equally impaired. Only the complemented strain and an overexpression strain showed measurable growth. These data suggest that the complemented strain is producing higher levels of *pe22/ppe36* than WT, and that while deletion reduces sulfate intracellularly, overexpression increases levels of sulfate transport. To assess the specificity of the PE22/PPE36 complex for sulfate, we next tested growth on phosphate-depleted medium by spotting experiments (Fig. 2B). No significant growth differences were observed among the wild-type, KO, and complemented strains across the same concentration range, suggesting that PE22/PPE36 is specific for sulfate.

**Fig 2.**
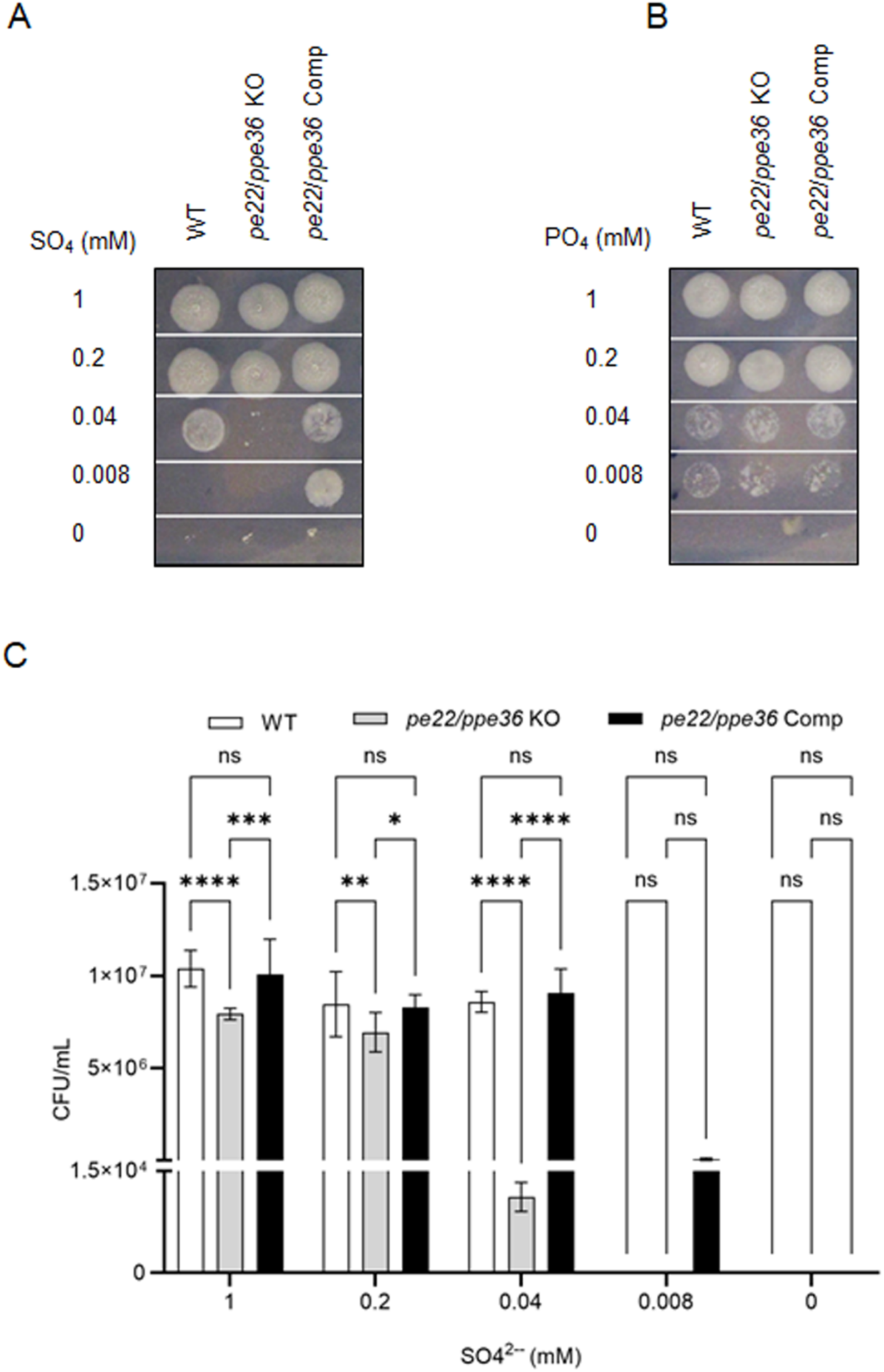
Loss of PE22/PPE36 impairs growth under sulfate-limiting conditions. Spotting of WT, *pe22/ppe36 KO*, and *pe22/ppe36* complemented (Comp) strains grown in (A) sulfate-free and (B) phosphate-free media, followed by repletion with gradient concentrations (1 mM, 0.2 mM, 0.04 mM, 0.008 mM, and 0 mM) of sodium sulfate or potassium phosphate, respectively, after 48 h of starvation. After 21 days of growth, all strains were spotted (3 µL) on 7H10 agar plates, and images were captured after 10 days of incubation. One representative image from three biological replicates is shown. (C) CFU enumeration of WT, *pe22/ppe36* KO, *pe22/ppe36* Comp, and *pe22/ppe36* OE strains after 21 days of repletion with gradient concentrations of sodium sulfate. All experiments were performed in three biological replicates. Significance was determined using two-way ANOVA comparing WT with pe22/ppe36 KO, Comp, and OE strains (*P < 0.05, **P < 0.01, ***P < 0.001, ****P < 0.0001).

### Loss of PE22/PPE36 reduces cellular sulfate content

To directly test the idea that PE22/PPE36 has a role in sulfate transport, we measured the intracellular concentration of elemental sulfur in WT, KO, and Comp strains. To measure total cellular elemental sulfur, we used Inductively Coupled Plasma Optical Emission Spectroscopy (ICP-OES) that uses a high-temperature argon plasma to atomize and excite a liquid sample. Different elements can then be detected and quantified based on their emission spectra. We normalized sample quantity by optical density and measured the intracellular sulfur at 0, 5min, 6h, and 24h. Because ICP-OES measures the total elemental sulfur pool in the cell, including sulfur from protein, we expected only modest changes of *pe22/ppe36* deletion in this assay. The *pe22/ppe36* KO showed ∼30% less total sulfur at 24 and 48h after repletion of media with sulfate (Fig. 3 A and B). The complemented strain recovered sulfate content to the level of WT. As a selectivity control, we also tested the uptake of the oxyanion analog phosphate in the *pe22/ppe36* KO but did not find any significant difference in phosphate levels (Fig 3 C), indicating specificity of PE22/PPE36 for sulfate.

**Fig 3.**
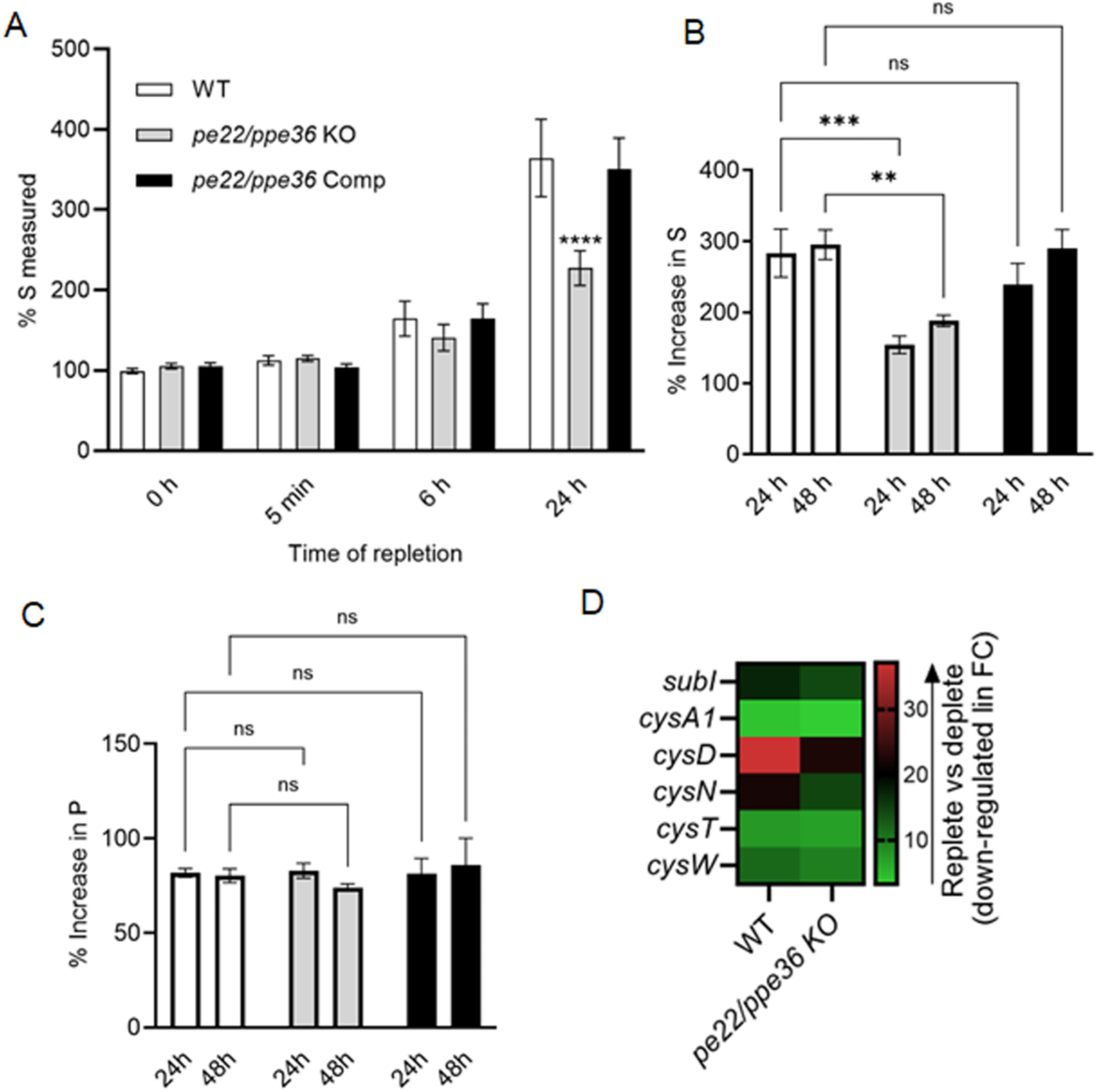
Loss of PE22/PPE36 specifically impairs sulfate uptake. (A) Sulfate uptake was measured in WT, *pe22/ppe36* KO, and *pe22/ppe36* Comp strains by ICP-OES after 48 h of starvation and repletion with 10 mM sodium sulfate at different time points (0 h, 5 min, 6 h, and 24 h). (B) Percent increase in intracellular sulfate was measured at 24 h and 48 h relative to 0 h after sulfate repletion. (C) Percent increase in intracellular phosphate was measured at 24 h and 48 h relative to 0 h under the same conditions. (D) Linear fold change showing the downregulation of sulfur-responsive genes *cysT, cysW, cysA2,* and *cysA3* in WT and *pe22/ppe36* KO strains based on RNA-seq data comparing sulfate-replete vs sulfate-depleted conditions. All experiments were performed in three biological replicates. Significance was determined using two-way ANOVA comparing WT with *pe22/ppe36* KO, Comp, and OE strains (*P < 0.05, **P < 0.01, ***P < 0.001, ****P < 0.0001).

### The transcriptional response to sulfate is muted in the *pe22/ppe36* KO strain

We next tested the transcriptional response to sulfate in the *pe22/ppe36* KO strain and WT by RNA-seq as a measure of intracellular sulfate. The known sulfur-responsive genes *cysT, cysW, cysA2,* and *cysA3* were downregulated in the WT, as previously noted, but less so in the *pe22/ppe36* KO strain, consistent with less intracellular sulfate in these cells. These data indicate that changes in PE22/PPE36 are sufficient to alter the intracellular sulfate levels (Fig. 3D).

### MspA complements *pe22/ppe36* KO

To test if the deletion of *pe22/ppe36* can be functionally complemented by a heterologous outer membrane porin, we expressed the major porin MspA from *Mycobacterium smegmatis* (*Msm*) in the *Mtb pe22/ppe36* KO background (*pe22/ppe36* KO+MspA) and measured intracellular sulfur by ICP-OES. MspA expression in the KO background reverted the loss of intracellular sulfate back to WT levels (Fig 4A), suggesting that an outer membrane porin can functionally complement the loss of *pe22/ppe36*.

**Fig. 4.**
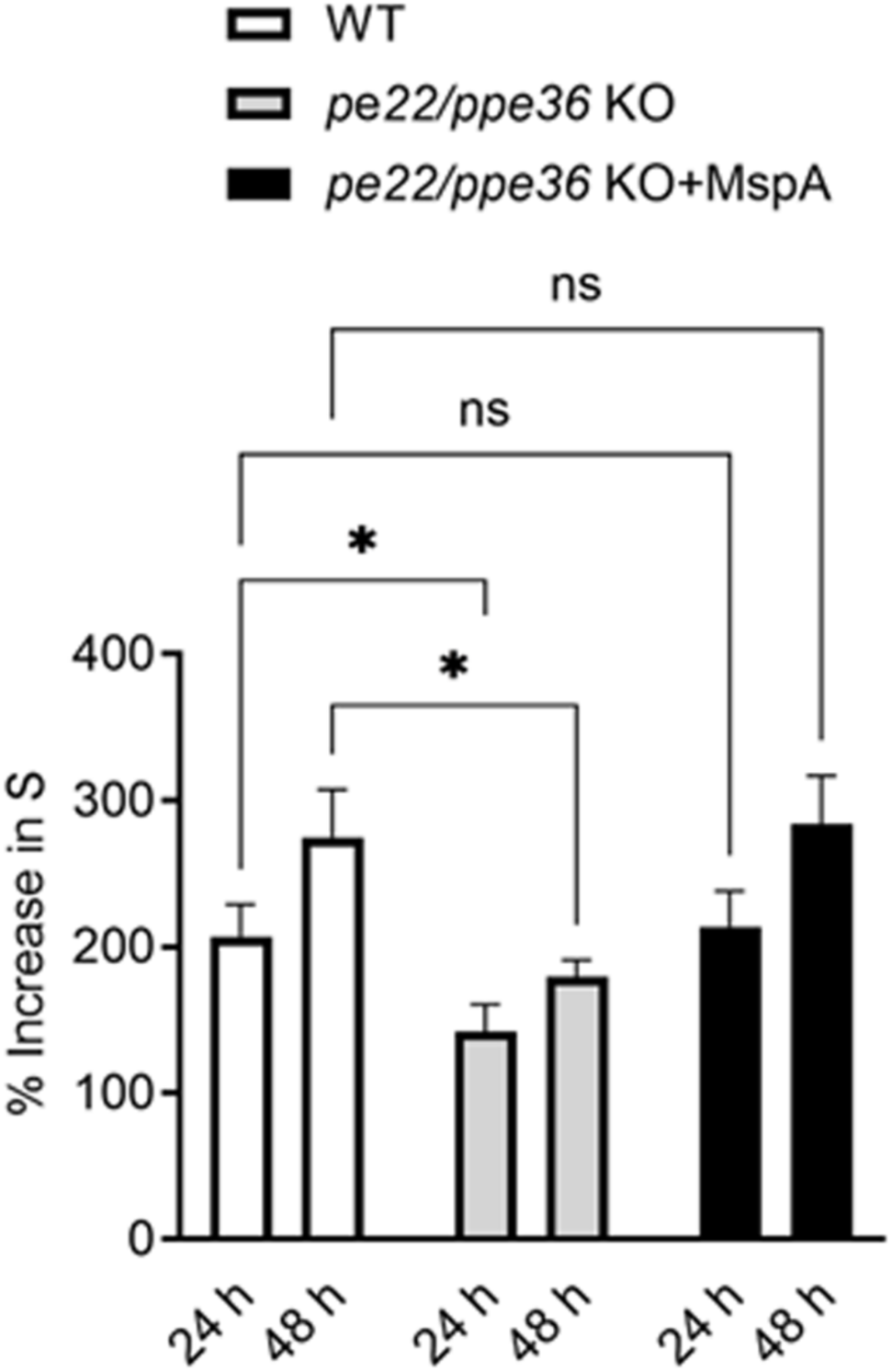
MspA complements the *pe22/ppe36* KO phenotype. (A) Percent increase in sulfate was measured in WT, *pe22/ppe36* KO, and *pe22/ppe36* KO + MspA strains at 24 h and 48 h relative to 0 h after 48 h of sulfate starvation and subsequent repletion. Significance was calculated relative to WT at each time point using two-way ANOVA (*P < 0.05, **P < 0.01, ***P < 0.001, ****P < 0.0001).

### Sulfolipid and sulfur metabolites are affected by *pe22/ppe36* KO

As a metabolic indicator of differences in cellular sulfate levels, we next tested for changes in sulfolipids in the OM as a result of *pe22/ppe36* deletion. We extracted total lipids from WT, KO and Comp strains and resolved lipids in a Chloroform:Methanol:Water solvent system by thin layer chromatography. The *pe22/ppe36* KO strain had less sulfolipids compared to WT (Fig. 5A and 5B). Complementation with episomal *pe22/ppe36* restored sulfolipid levels to WT, suggesting that *pe22/ppe36* KO affects downstream sulfolipid levels in the cell wall. However, TLC did not resolve individual sulfolipid and other lipid species, and SL-1 and phthiocerol dimycocerosates (PDIM) production, for example, are linked in *Mtb*. To further parse the effects of *pe22/ppe36* deletion on downstream metabolites and sulfolipids, we next measured sulfate-containing metabolite and lipid levels in the WT, KO, and complemented strains by mass spectrometry (MS).

**Fig. 5.**
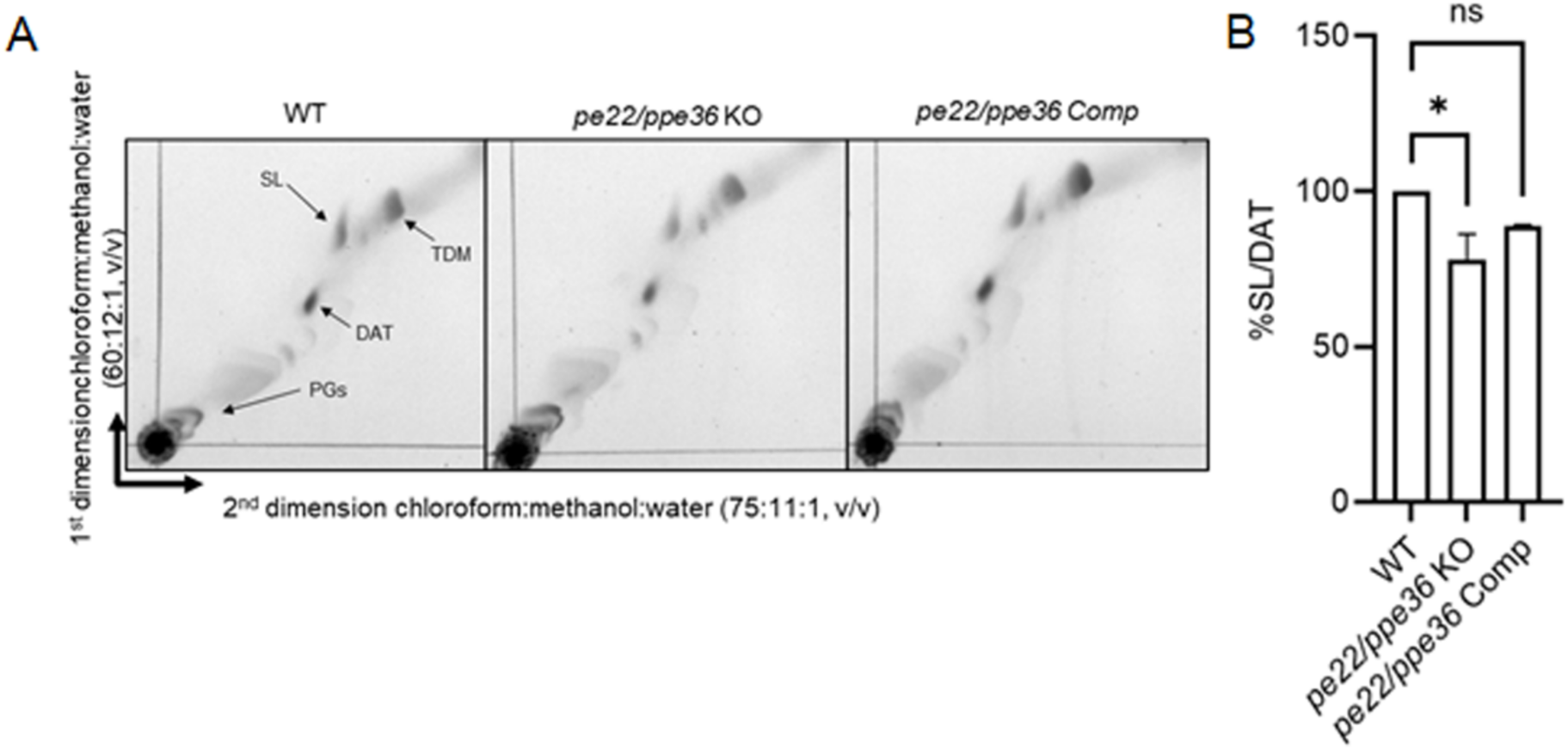
PE22/PPE36 affects levels of cell wall sulfolipid. PE22/PPE36 affects cell wall sulfolipid (SL) levels. (A) Total lipids were extracted from WT, *pe22/ppe36* KO, and *pe22/ppe36* Comp strains and resolved by two dimensional thin-layer chromatography using a chloroform:methanol:water solvent system in (75:11:1) and (60:12:1) ratio for first and second dimension, respectively. (B) Intensity of sulfolipid normalized to the non-sulfur–containing lipid diacyltrehalose (DAT) from the same TLC plate, plotted as the percentage of SL/DAT intensity for each strain based on two independent experiments.

Once imported, sulfate is activated by ATP sulfurylase (*cysND*) to form the metabolic branchpoint intermediate, adenosine-5′-phosphosulfate (APS) (11, 22, 23). APS can be phosphorylated by APS kinase (*cysC*) to produce PAPS, the universal sulfate donor for sulfotransferases (STs) (11, 23, 24). STs transfer sulfate from PAPS to hydroxyl or amide groups on glycolipids to produce sulfolipids, which have central roles in cell wall integrity (25–27). We tested if the sulfate import deficits of the *pe22/ppe36* deletion strain impacted levels of downstream lipid products in the presence of exogenous sulfate. These studies showed a selective reduction in levels of mature tetra-acylated sulfolipids that were matched by a reciprocal increase in levels of PDIM, both of which compete for a shared pool of the methyl branched lipid precursor, methylmalonyl-CoA (Fig. 6). The sulfolipid defects could be genetically complemented by expression of an ectopic copy of *pe22/ppe36* or *mspA*. In contrast to sulfolipids, PDIM levels in the MspA-expressing strain were not reduced to the same extent as in the complemented strain (Fig. 6, lower panel). This difference could reflect the more non-specific transport properties of MspA and an effect on uptake of other medium components such as propionate, which can impact methyl-branched lipid biosynthesis. Levels of APS-derived metabolites, in contrast, exhibited no significant change, suggesting that they are quickly shunted toward *de novo* lipid synthesis under conditions supporting active replication (data not shown).

**Fig. 6.**
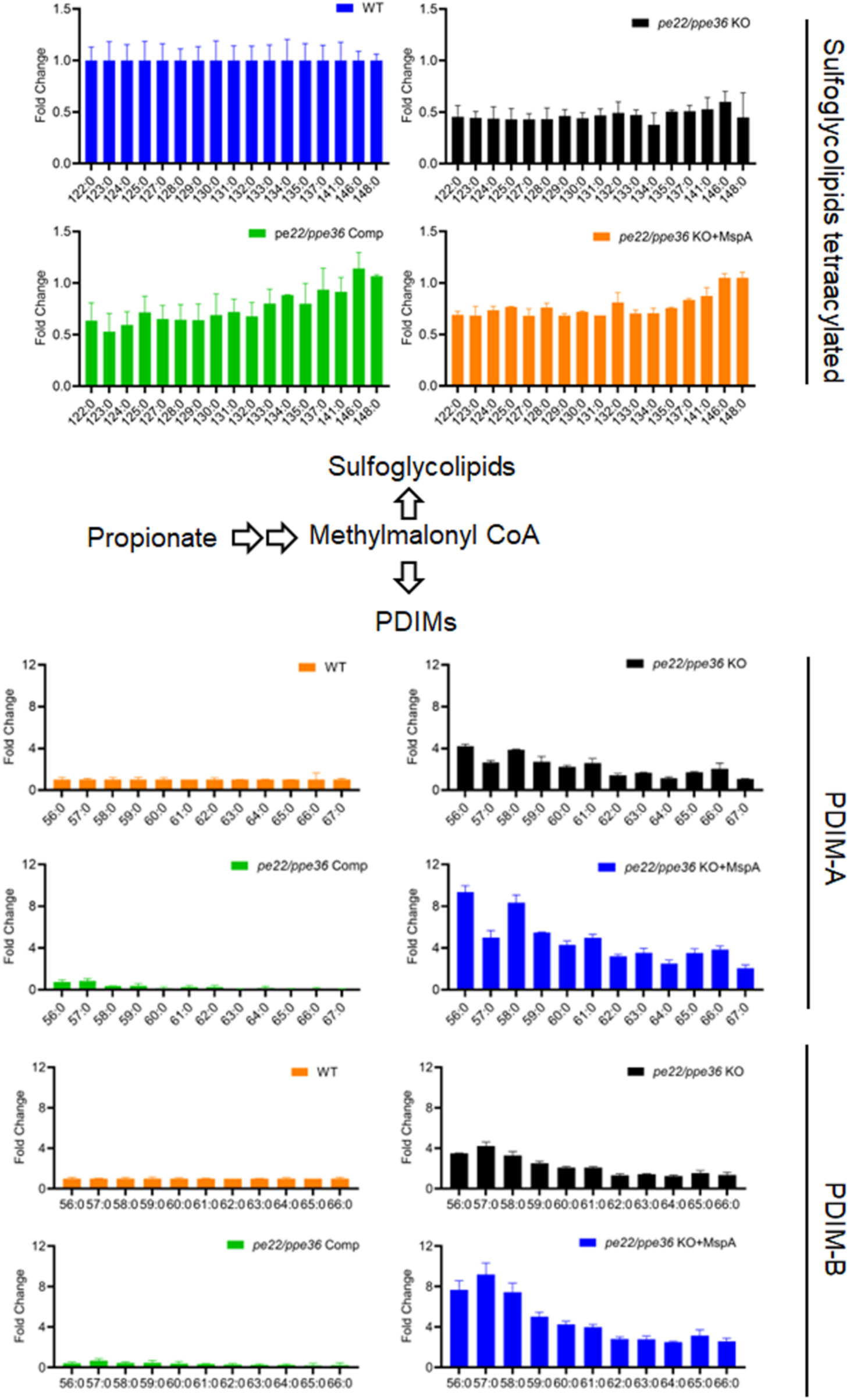
Sulfate import via PE22/PPE36 selectively influences sulfolipid and PDIM abundance. LC–MS/MS–based lipidomic analysis of total lipid extracts from WT, *pe22/ppe36* knockout KO, and *pe22/ppe36* Comp strains. Shown are the relative abundances of sulfoglycolipids (top panel) and PDIM-A and PDIM-B species (bottom panel). For the sulfoglycolipid bar graph, the x-axis indicates the total carbon chain length of tetra-acylated species, with “0” denoting unsaturated acyl chains. For PDIMs, the numbers indicate the total carbon chain length of the phthiocerol dimycocerosate species. Average of two independent experiments is shown.

### *pe22/ppe36* deletion does not affect general OM permeability

Deletion of *pe22/ppe36* affected PDIM levels in the outer membrane (Fig 6). PDIM can also alter outer membrane permeability. To test if the differences in sulfate levels in the *pe22/ppe36* KO are due to compromised import, or an indirect effect of an increase in PDIM, we tested the general permeability of the *pe22/ppe36* KO by an EtBr uptake assay. We did not detect a significant difference between the WT and *pe22/ppe36* KO strains in EtBr uptake (Fig. S1). If anything, the *pe22/ppe36* KO, which had higher PDIM levels, allowed marginally more EtBr into the cell, an effect consistent with higher permeability, not less. In addition, we showed by ICP-OES that levels of the chemically related oxyanion phosphate were not altered in the *pe22/ppe36* KO. We also recently showed that the deletion of *pe22/ppe36* did not affect the susceptibility to any of ten clinical tuberculosis drugs, suggesting that higher PDIM in the deletion strain does not affect the outer membrane permeability (6). These data show that elevated PDIM levels in the *pe22/ppe36* KO are a secondary effect, not the driver of reduced sulfate uptake.

## Discussion

Due to their large footprint in the *Mtb* genome and their association with pathogenic mycobacteria, the PE/PPE proteins have been the object of much speculation. Twenty-five years after they were first described, a shared role among at least some members of this diverse protein family is emerging: Several recent studies have shown functions in outer membrane transport for multiple PE/PPEs across different subfamilies. Consistent with other bacterial membrane transporters, PE/PPEs have now also been shown to inadvertently transport drugs. These data also offer reinterpretations of previous findings that implicated PE/PPE proteins in acquisition of nutrients that are also consistent with the transporter paradigm. Given the large number of PE and PPE proteins, their unusual, partially repetitive sequences and the almost complete lack of structural data, much remains to be understood before the full extent and consequence of the PE/PPEs for outer membrane transport will be clearly defined. But the current findings together point to a remarkable and surprising solution to one of the central questions of mycobacterial physiology, outer membrane transport, that has specifically evolved in pathogenic mycobacteria and bears little resemblance to other known bacterial or eukaryotic systems.

Part of the reason why the PE/PPEs have so long remained intractable is that working with them is challenging on every level: DNA sequencing, in particular short read-based sequencing is complicated by the long repeat regions, mass spectrometry-based proteomic analyses are complicated by the low numbers of canonical protease cleavage sites, recombinant expression is challenging because of the requirements for specialized secretion systems, and genetic analysis is complicated by the likely redundancy between the many paralogs in the family. Because redundancy is likely among the *pe/ppe* genes, we expected only modest phenotypes of our *pe22/ppe36* deletion strain. However, the growth defects in sulfate-depleted media were clear, and the reduction of total elemental sulfur in the deletion strain as measured by ICP-OES was markedly reduced, suggesting that *pe22/ppe36* facilitates a large part of sulfate uptake.

Somewhat surprisingly, the PE22/PPE36 complex was previously shown to be involved in heme acquisition (5). PE22/PPE36 appears to be specific for sulfate over other oxyanions, so how would it transport such different chemical entities such as sulfate and heme? Structural data on the particular substrate binding site(s) and transmembrane regions will help in resolving these questions. Promiscuous transport activity is not unique to PE22/PPE36 but has also been shown in a recent study that showed drug import of several PE/PPEs (6).

Our data show an effect of *pe22/ppe36* deletion on intracellular sulfate on several levels, including a reduced content of sulfolipids accompanied by a reciprocal increase in PDIMs. The metabolic coupling of the two through the shared precursor methylmalonyl-CoA has previously been established (28), and with reduced flux of methylmalonyl-CoA through one pathway due to lack of sulfate, a seesaw to the other is expected. These key components of the outer membrane were long thought to be associated with virulence of *Mtb*, but more recent studies suggest they are not. Sulfolipids do appear to play a role in pathogenicity, however; they have been shown to affect coughing (29), the central route of *Mtb* transmission. These data show how perturbation of *pe/ppe* genes can lead to pathogenicity phenotypes, but that those are secondary to the primary function in nutrient transport. These data add a cautionary note to the frequent characterization of the *pe/ppe* genes as virulence factors; even if virulence phenotypes in *pe/ppe* mutant strains are observed, these may not always reflect a specifically evolved virulence function but a collateral effect of a nutrient imbalance.

The source of sulfate for *Mtb in vivo* was long unclear and inorganic sulfate has only recently been shown to be the main sulfur source *in vivo*. In addition to further supporting the emerging PE/PPE transporter paradigm, we here show the mechanism by which inorganic sulfur can cross the outer membrane—a remarkable structure that requires not only unique chemistry but also unique transport processes.

## MATERIALS AND METHODS

### Media and growth conditions

We used *Mtb* H37Rv as the parental strain for generating all mutants. Strains were grown in Middlebrook 7H9 medium (Difco) supplemented with 10% (v/v) oleic acid albumin dextrose catalase (OADC) enrichment (BBL; Becton Dickinson), 0.5% glycerol, 0.02% Tyloxapol and 100 µM sodium propionate. Media were designated as follows: 7H9+GOTy (Tyloxapol), and 7H9+GOTyP (Tyloxapol plus 100 µM sodium propionate) to maintain PDIM levels (30). For solid culture, Middlebrook 7H10 agar was supplemented with 10% OADC and 0.5% glycerol. For sulfate starvation, modified sulfate-free Sauton’s medium was used, containing L-asparagine (4 g/L), citric acid (2 g/L), ferric ammonium citrate (0.05 g/L), MgCl₂ (0.5 g/L), K₂HPO₄ (0.5 g/L), ZnCl₂ (0.1 mg/L), and glycerol (2% v/v), in which all sulfate salts were replaced with the corresponding chloride salts. Phosphate-free medium was prepared with KCl (0.27 g/L), MgSO₄ (0.5 g/L), L-asparagine (4 g/L), citric acid (2 g/L), ferric ammonium citrate (0.05 g/L), ZnSO₄ (0.1 mg/L), and glycerol (2% v/v). For both sulfate- and phosphate-free media, the pH was adjusted to 7.0, followed by the addition of 0.02% Tyloxapol and 100 µM sodium propionate, and the final volume was adjusted to 1 L. Strains harboring antibiotic resistance cassettes were cultured with the appropriate antibiotics: hygromycin (50 µg/mL), kanamycin (25 µg/mL), or zeocin (25 µg/mL). All drugs used for screening assays were purchased from Sigma, and stock solutions were prepared in either water or DMSO.

### Creation of deletion and complemented strains

The deletion (KO) strains were generated using recombineering, as described previously (19). Initially, 500 bp upstream and downstream of the gene of interest, along with the hygromycin resistance cassette (primers Hyg_F and Hyg_R) or zeocin resistance cassette (primers Zeo_F and Zeo_R), were amplified separately (all primers used for each deletion are listed in Supplemental Table S1). The PCR fragments were then Gibson ligated to the 5′ and 3′ ends of the hygromycin or zeocin resistance cassette to create the recombineering cassette. This linear recombineering cassette was PCR-amplified, purified, and electroporated into the *Mtb* H37Rv strain carrying the recombineering plasmid pNIT:Etc (31). Hygromycin-resistant colonies were screened, and the positive clones were confirmed by DNA sequencing.

For the complemented strains, the *pe22/ppe36,* coding regions, as well as the *M. smegmatis mspA* coding region were amplified by PCR using primers listed in Supplemental Table S1 and Gibson cloned into the pDTCF plasmid (Zeocin resistance) with a C-terminal FLAG tag, under the control of an anhydrotetracycline (ATc)-inducible promoter. The resulting plasmids were electroporated into the H37Rv *pe22/ppe36* deletion strain, and positive clones were selected by growth in hygromycin and zeocin.

### RNA sequencing

The H37Rv (WT) and *pe22/ppe36* KO strains were grown to an OD_600_ of 0.8 in 7H9+GOTyP medium. For RNA-seq of WT H37Rv in with and without sulfate media cells were washed three times to remove any trace of sulfate from 7H9 media and then grown in sulfate-free HdB media with 100 µM sodium sulfate or without sulfate for further 48 h. Cells were pelleted at 4,000 g for 5 minutes at 4°C, resuspended in Trizol, and lysed by bead beating for 30 seconds at 6 m/s for 3 cycles, with intermittent cooling on ice. Cell debris was pelleted at 20,000 g for 1 minute, and the supernatant was transferred to a heavy phase-lock gel tube containing 300 µL of chloroform. The tubes were inverted for 2 minutes and centrifuged at 20,000 g for 5 minutes. RNA in the aqueous phase was precipitated with 300 µL of isopropanol and 300 µL of high-salt solution (0.8 M sodium citrate, 1.2 M sodium chloride). RNA was purified using the QIAGEN RNeasy kit, and ribosomal RNA was depleted using a previously published protocol (32). Briefly, a biotinylated oligo mixture of 23S, 16S, and 5S was incubated with RNA to anneal to rRNA, followed by incubation with streptavidin beads. mRNA was purified from the supernatant using Ampure XP beads. A cDNA library was generated using the NEBNext Ultra II RNA Library Prep Kit, and each replicate was barcoded in the DNA library using the NEBNext Multiplex Oligos for Illumina. Libraries were quantified using the KAPA qPCR quantification kit, pooled, and sequenced at the University of Washington Northwest Genomics Center using the Illumina NextSeq 500 High Output v2 Kit.Read alignment was performed using the Bowtie 2 custom processing pipeline (https://github.com/robertdouglasmorrison/DuffyNGS, https://github.com/robertdouglasmorrison/DuffyTools). Gene expression changes were identified using a combination of five differential expression tools within DuffyTools: round robin, RankProduct, significance analysis of microarrays (SAMs), EdgeR, and DeSeq2. The results from each DE tool were combined using a weighted average of fold change and significance (*P-*value). Genes with an averaged absolute fold change of more than 2-fold and a *P-*value <0.01 were considered differentially expressed.

### Culture condition for sulfate repletion and ICP-OES

H37Rv (WT), the *pe22/ppe36* knockout (*pe22/ppe36* KO) and complemented (*pe22/ppe36* Comp) strains were used for sulfate repletion experiments. Strains were routinely cultured in Middlebrook 7H9 broth supplemented with 0.2% (v/v) glycerol, 10% (v/v) OADC enrichment, 0.02% (v/v) tyloxapol, and 100 µM sodium propionate. Cultures were grown at 37°C with gentle agitation until mid-log phase (OD₆₀₀ ≈ 0.6–0.8). Where indicated, gene expression was induced by addition of anhydrotetracycline (ATc) for at least 48 h prior to sulfur depletion. All experiments involving live *Mtb* were performed under BSL-3 containment following institutional biosafety guidelines.

Mid-log-phase cultures were harvested by centrifugation and washed three times with sterile phosphate-buffered saline (PBS) to remove residual sulfur from the growth medium. Washed cells were resuspended in sulfate-free Sauton’s medium supplemented with 0.02% tyloxapol, 100 µM sodium propionate, and ATc (prepared in DMF to avoid sulfur contamination). Cultures were incubated for 48 h at 37°C to deplete intracellular sulfur pools. Following sulfur starvation, cultures were normalized to equivalent OD₆₀₀ values and split into sulfate-replete and sulfate-deplete conditions. Sulfate repletion was initiated by addition of sodium sulfate (Na₂SO₄) to a final concentration of 10 mM, while control cultures received no sulfate. Cultures were incubated for an additional 24 or 48 h prior to harvesting, depending on the downstream analysis. For elemental analysis following sulfate repletion, three OD-equivalent cultures per condition were harvested by centrifugation at 4,000 × g for 5 min at 4°C. Cell pellets were washed twice with Chelex-treated Milli-Q water containing 0.02% tyloxapol to remove extracellular ions. After the final wash, pellets were resuspended in 2.8 mL Chelex-treated water and aliquoted into three 950-µL O-ring screw-cap tubes. Samples were heat-inactivated by boiling at 110°C for 30 min with intermittent inversion.

Nitric acid was added to each tube to a final concentration of approximately 2% (v/v), and samples were incubated overnight at 75°C to ensure complete digestion. The following day, triplicate digests from the same sample were pooled in 15 mL tubes under a chemical fume hood. Elemental standards were prepared in 0.1% nitric acid, and serial dilutions were generated when required. Samples were analyzed by inductively coupled plasma–optical emission spectrometry (ICP-OES) according to standard operating procedures, and sulfur content was quantified relative to appropriate calibration standards and media controls.

### Total Lipid Extraction

For lipid analysis, *Mtb* H37Rv and KO and Comp strains were grown in sulfate-free Sauton’s medium and subjected to sulfate repletion for 72 h as described above. Fifty milliliters of culture at an OD₆₀₀ of approximately 1.0 were harvested by centrifugation, and cell pellets were washed three times with phosphate-buffered saline (PBS) to remove residual media components. Washed pellets were stored at −80°C until lipid extraction.

At the time of extraction, frozen pellets were resuspended in 2 mL of methanol and transferred to 15-mL glass conical tubes fitted with Teflon-lined caps. Chloroform (4 mL) was added using glass pipettes to obtain a final solvent ratio of 2:1 (chloroform:methanol). Tubes were protected from light and incubated overnight with gentle shaking at room temperature inside a biosafety cabinet to allow lipid extraction. The following day, tubes were placed upright to allow cellular debris to settle by gravity, and the supernatant was carefully transferred to a fresh glass tube. The supernatant fractions were sterile-filtered sequentially using a glass syringe through one 0.4-µm PTFE filter followed by two 0.2-µm PTFE filters. The tube with filtered lipid extracts were surface-decontaminated with Oxivir wipes (three treatments) prior to removal from BSL-3 containment, ensuring that all tubes were clearly labeled due to potential label loss during solvent handling. Filtered lipid extracts were transferred in 1-mL aliquots into pre-weighed amber glass vials and dried under a continuous stream of nitrogen gas using a heat block at 37°C. Solvent evaporation was performed gradually by repeatedly adding extract aliquots until all solvent was removed. The mass of extracted lipid was determined gravimetrically by subtracting the weight of the empty vial. Dried lipid samples were stored at −80°C until further analysis.

### Thin-Layer Chromatography (TLC) Analysis

For TLC analysis, dried lipid extracts were resuspended in chloroform:methanol (2:1, v/v) to a final concentration of 20 mg/mL. Total lipids (300 µg per lane; 15 µL) were spotted onto silica gel TLC plates and resolved using petroleum ether:ethyl acetate (98:2, v/v) as the mobile phase, unless otherwise indicated. After solvent migration, plates were air-dried and sprayed with 5% (v/v) sulfuric acid in methanol. Plates were then heated using a hot air blower until lipid bands developed and became visible. TLC profiles were documented and compared across sulfate-replete and - deplete conditions to assess changes in lipid composition.

### HPLC-coupled electrospray ionization lipidomic profiling

Lipidomic datasets were generated using a 1600 series HPLC system with an inline Varian Monochrom diol guard column (3 μm x 4.6 mm) and Varian Monochrom diol column (3 μm x 150 mm x 2 mm), coupled to an Agilent Technologies 6545 Accurate-Mass Q-Tof mass detector. Dried lipid extracts were resuspended at 0.5 mg/mL in solvent A [hexanes: isopropanol 70:30 (v: v), 0.02% (m:v) formic acid, 0.01% (m:v) ammonium hydroxide], filtered or centrifuged at 1500 rpm for 5 min to remove trace non-lipidic materials prior to transfer to a glass autosampler vial (Agilent). Ten μg of lipid extracts were injected and chromatographed at 20 C and flow rate of 0.15 mL/min with a binary gradient from 0% to 100% solvent B [isopropanol: methanol 70:30 (v:v), 0.02% (m:v) formic acid, 0.01% (m:v) ammonium hydroxide]: 0-10 min, 0% B; 17-22 min, 50% B; 30-35 min, 100% B; 40-44 min, 0% B, followed by additional 6 min 0% B post-run. Ionization was maintained at 325°C with a 5L/min drying gas flow, a 30 psi nebulizer pressure and 5500 volts. Spectra were collected in positive and negative-ion mode from *m/z* 100 to 3000 at 1 spectrum/s. Continuous infusion calibrants included *m/z* 121.050873 and 922.009798 in positive-ion mode and *m/z* 112.985587 and 1033.98810 in negative-ion mode.

### Accumulation of Ethidium Bromide

WT, *pe22/ppe36* KO, *pe22/ppe36* Comp and *pe22/ppe36* KO +MspA strains were grown in Middlebrook 7H9 medium supplemented with OADC at 37 °C until mid-log phase (OD₆₀₀ = 0.6–0.8). Cultures were harvested by centrifugation at 2,880 × g for 10 min, washed once, and resuspended in phosphate-buffered saline (PBS; pH 7.4). The bacterial suspension was adjusted to an OD₆₀₀ of 0.8 in PBS containing 0.05% Tween 80 and 0.4% (w/v) glucose. Aliquots of 100 µL were dispensed into black 96-well microplates with transparent bottom and mixed with an equal volume of ethidium bromide (EtBr) solution (1 µg/mL), resulting in a final OD₆₀₀ of 0.4 and a final EtBr concentration of 0.5 µg/mL. Intracellular accumulation of EtBr was monitored at 37 °C for 60 min using a BMG PHERAstar plate reader, with excitation at 540 ± 20 nm and emission at 590 ± 20 nm. Fluorescence readings were acquired at every 1 min till 60 min of total time. Two controls were included in each experiment: (i) EtBr-only control, consisting of 100 µL PBS mixed with 100 µL EtBr without bacterial cells, to determine background fluorescence; and (ii) cell-only control, consisting of 100 µL PBS mixed with 100 µL bacterial suspension without EtBr, to account for intrinsic cellular fluorescence. Relative fluorescence values were obtained after subtraction of background fluorescence.

---

**Fig. S1.**
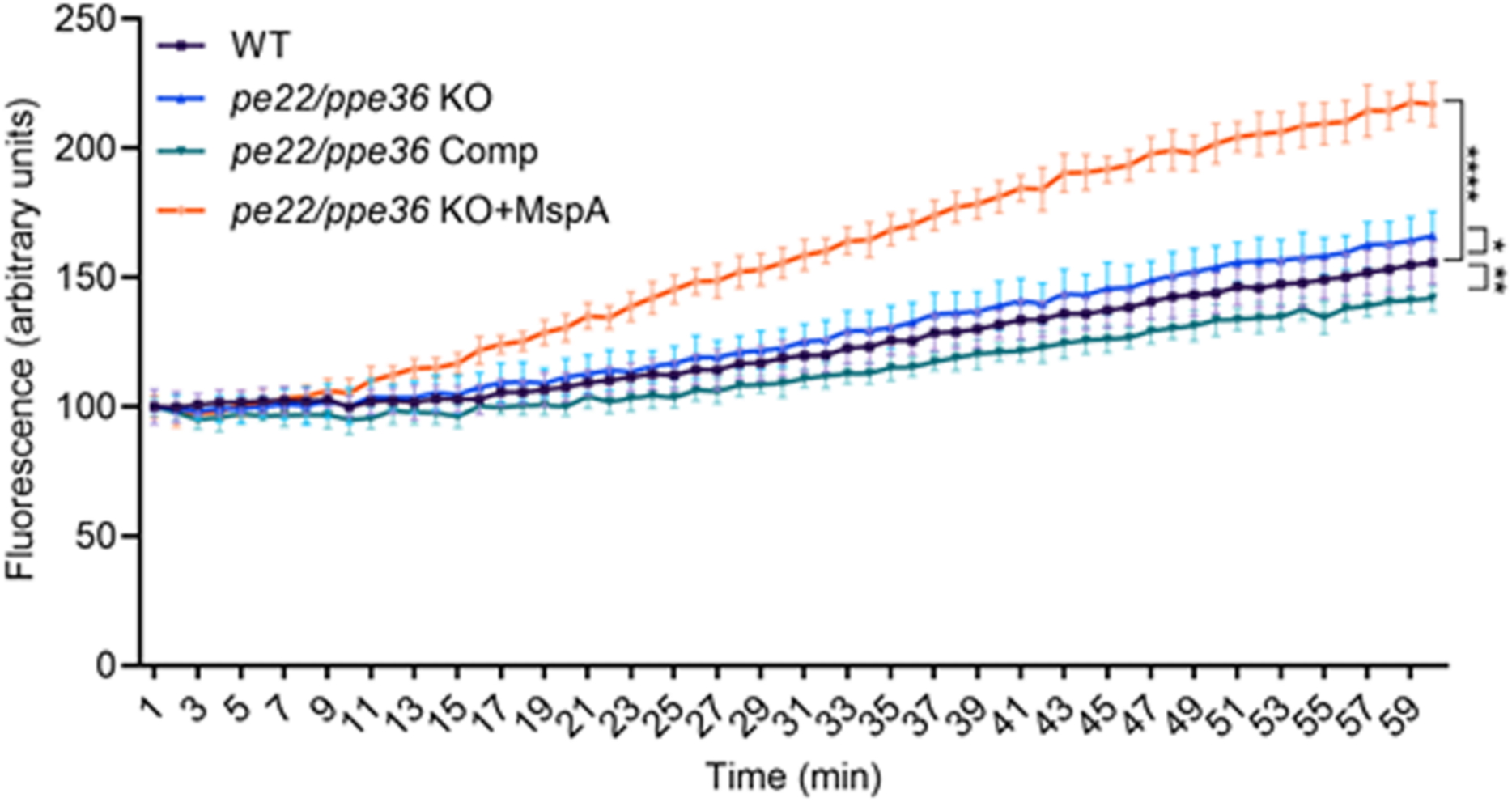
Depletion of PE22/PPE36 does not affect general OM permeability. EtBr uptake assay was performed on all strains in exponential phase. Significance was calculated relative to WT at each time point using two-way ANOVA (*P < 0.05, **P < 0.01, ***P < 0.001, ****P < 0.0001).

**Table S1:**
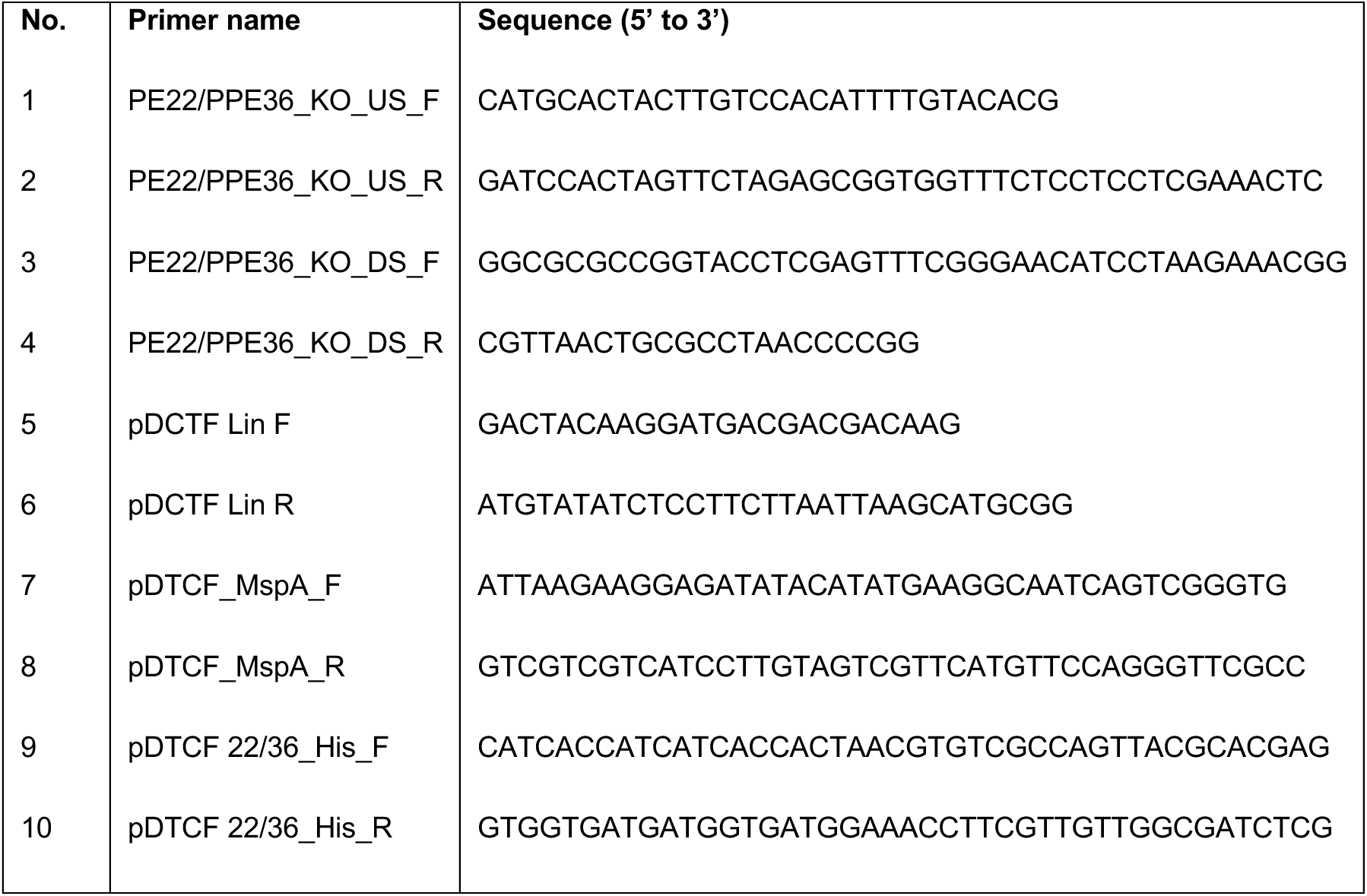
List of primers used in this study.

